# A StayGold-based calcium ion indicator

**DOI:** 10.64898/2026.03.06.710044

**Authors:** Ikumi Miyazaki, Kelvin K. Tsao, Takuya Terai, Kei Takahashi-Yamashiro, Robert E. Campbell

## Abstract

Genetically encoded calcium ion (Ca^2+^) indicators (GECIs) enable visualization of Ca^2+^ dynamics in living systems but often suffer from limited photostability during prolonged imaging. The recent discovery of StayGold, a green fluorescent protein (FP) with exceptional brightness and photostability, opened the possibility of addressing this longstanding challenge. Here, we sought to establish whether a monomeric variant of StayGold (mStayGold) could be converted into a single FP-based GECI. Through extensive protein engineering, we generated a functional mStayGold-based GECI, HiCaRI (Highly intensiometric Ca^2+^ Responsive Indicator) by fusing Calmodulin (CaM) and the ckkap binding peptide from K-GECO1 into mStayGold(J). HiCaRI exhibits a large Ca^2+^-dependent inverse fluorescence response (Δ*F*/*F*_min_ = −15) while retaining high brightness and improved photostability relative to previously reported GFP-based GECIs. Although the current variant represents a first-generation prototype with shortcomings in terms of Ca^2+^ affinity and photostability (relative to StayGold and mStayGold(J)), this work demonstrates the feasibility of constructing single FP-based GECIs from a highly photostable fluorescent protein.

## Introduction

The discovery and subsequent engineering of *Aequorea victoria* green fluorescent protein (GFP) laid the foundation for the development of genetically encoded calcium ion (Ca^2+^) indicators (GECIs), that have become essential tools for visualizing intracellular Ca^2+^ dynamics with high spatial and temporal resolution.^1–3^ Baird et al. developed the first single FP-based Ca^2+^ indicator, Camgaroo1, by inserting calmodulin (CaM) into yellow FP (YFP) such that Ca^2+^-binding induced a conformational change that altered fluorescence.^4^ In the same work, Baird et al. reported circularly permutated (cp) YFP variants and proposed a variety of topologies for insertion of other proteins into GFP or insertion of cpGFP into other proteins. The strategy of cpGFP insertion was later adopted for the creation of the cpGFP-based Pericam and G-CaMP GECIs, demonstrating the feasibility of coupling the Ca^2+^-dependent conformational change of a CaM plus a CaM-binding peptide to substantial changes in fluorescence.^5,6^ Although early generations exhibited limited stability, folding, and brightness, iterative optimization through directed evolution and rational design produced increasingly bright and fast variants derived from G-CaMP. Recent examples include GCaMP6, jGCaMP7, and jGCaMP8, which enable sensitive detection of neuronal signals and action potentials and enjoy widespread use as GECIs for cellular and *in vivo* Ca^2+^ imaging.^3,7–9^ However, despite these advances, the current GCaMP derivatives and other single FP-based GECIs all exhibit limited photostability arising from the intrinsic properties of the fluorescent protein scaffold itself. With continuous or repeated illumination, the FP undergoes substantial photobleaching, limiting its long-term and high-intensity imaging applications that require a stable baseline fluorescence over an extended period.^10,11^ Furthermore, in most fluorescent protein scaffolds, there is an intrinsic trade-off between brightness and photostability, hence it is challenging to engineer variants that improve both.^12–14^

In 2022, a new class of GFP called StayGold was discovered from *Cytaeis uchidae* and immediately gained attention for its exceptional brightness and photostability compared to *Aequorea* GFP variants, which have long served as the scaffold for biosensors.^14^ However, the original StayGold was dimeric, limiting its usage for protein tagging and biosensor development. In the vast majority of genetically encoded biosensors, sensing domains are inserted into the bulge region where the chromophore is closest to the protein surface, such that efficient coupling of ligand-induced conformational changes to chromophore fluorescence can be achieved.^15^ The bulge region also happens to be localized in the dimer interface. Since dimerization constrains these interfaces and limits design flexibility, monomerization is generally required before domain insertion for biosensor development.^16^

Motivated by the limitations imposed by its dimeric character, three research groups independently reported monomeric variants of StayGold within the two years following its introduction, each using a different approach. The mStayGold (E138D) was generated with a single structure-guided mutation in the dimer interface.^17^ In contrast, mStayGold(J) and mBaojin were derived through a combination of directed evolution and rational mutagenesis that targeted residues within the dimer interface.^18,19^ These three monomeric StayGold variants opened the possibility of developing new GECIs that could potentially exhibit improved photostability relative to previously reported GECIs. Indeed, Chelykhova et al. recently reported icBTnC2, a StayGold-based GECI.^20^ icBTnC2 is based on mBaoJin, a monomeric StayGold variant reported by the same lab in 2024,^19^ and was engineered by inserting the troponin C domain from the hummingbird *Calypte anna* into mBaoJin. icBTnC2 exhibits an inverse Ca^2+^ response and was optimized through directed evolution to improve Ca^2+^-dependent fluorescence change.

Here, we report the engineering and development of the first-generation mStayGold-based GECI (HiCaRI) from mStayGold(J) and the CaM/ckkap Ca^2+^ binding domain from K-GECO1, which exhibits brightness and kinetics that are sufficient for imaging of Ca^2+^ dynamics in mammalian cells.^21^ We chose to work with mStayGold(J), which strikes a balance between photostability and brightness.^18,22^ The current variant exhibits robust Ca^2+^-dependent inverse fluorescence changes (Δ*F*/*F*_min_ = −15) and reduced photobleaching compared to recent-generation GCaMP series GECIs. These results demonstrate the potential of using StayGold as a FP domain within the context of single FP-based biosensors, which may serve as a promising new platform for long-term Ca^2+^ imaging.

## Results

### Engineering an insertion-tolerant mStayGold scaffold for GECI development

Our strategy to develop a mStayGold-based GECI (mSG-GECI) was to construct a prototype by inserting the Ca^2+^ binding protein comprising CaM and a peptide, similar to how previous single FP-based GECIs were developed.^15,21^ First, a structural comparison with established GECIs was undertaken to guide the design of the mSG-GECI. Using the reported monomeric mStayGold(J) sequence and available crystal structures, we observed that the gatepost residues, serine (S143) and asparagine (N146), defined here as residues adjacent to the chromophore and closest to the protein surface resembles that of K-GECO1 in identity, position, and conformation (**Figure 1A**).^18,21^ Additionally, the bulge region, L144 and P145 residues having their side chains pointed towards the solvent, also resembles the bulge region of GFP.^15^ This similarity suggested that transplanting the linker-CaM-ckkap sensing domain from K-GECO1 onto non-cp mStayGold(J) or cp-mStayGold(J) could be a promising starting point. For the cpmSG-GECI design, we inserted additional C-terminal GFP-derived residues to match the length and composition of the jGCaMP8 linker.^9^ Gatepost residues, S143 and N146, were not randomized at this stage, as their polar character and structural positioning are likely critical for both chromophore protonation equilibria and the photostability of mStayGold(J).^18^ The resulting constructs, both the cp and non-cp, exhibited a complete loss of fluorescence and solubility compared to the original mStayGold(J) protein. The attempt to restore fluorescence via random mutagenesis and rational design was unsuccessful, suggesting that mStayGold(J) itself may not be intrinsically tolerant of domain insertion. We hypothesized that this may be due to prior structural studies showing that mStayGold(J) adopts a rigid, tightly packed β-barrel, in which insertions near the chromophore can readily disrupt folding and fluorescence.^19^ Therefore, we performed a systematic insertion-site screening to identify permissive regions for domain insertion in non-cp mStayGold(J). Specifically, a short, flexible linker (GGTGGSGG) was introduced at multiple positions within the β-strand loop closest to the chromophore (V142–E147), combined with zero-, one-, or two-residue deletions (**Figure 1B**). As a result, we found a variant (hereafter referred to as i-mSG-v.0.0) that retained weak but detectable fluorescence (less than 0.1% of mStayGold(J)). This suggested that the corresponding site could tolerate structural perturbation without complete loss of chromophore function, providing a viable starting point for engineering an insertion-tolerant mStayGold(J) variant. We then performed directed evolution on i-mSG-v.0.0, where we used error-prone PCR to introduce random mutations across the entire construct and select variants based on fluorescence brightness.^23^ After four rounds, we generated an insertion-tolerant i-mSG-v.1.0 with 7 mutations, with restored fluorescence and solubility that allowed us to move on to the GECI development (**Figure 1C, S1**).

**Figure 1.**
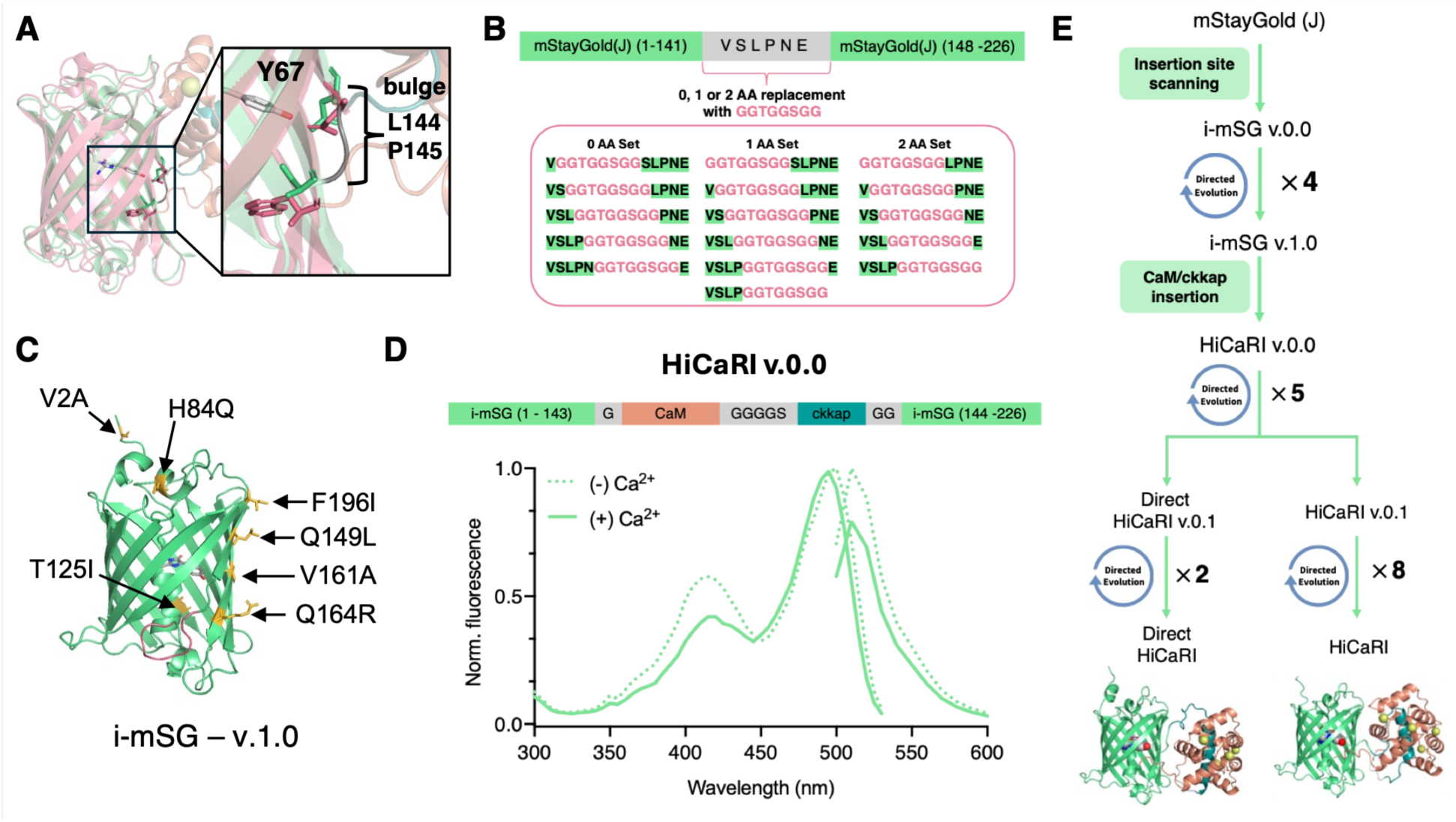
Overview of HiCaRI design and engineering. (**A**) Overlay of K-GECO1 (PDB: 5UKG) and mStayGold(J) (PDB: 8ILK).^14,21^ S and N gateposts of mStayGold(J) colored in green. W and N gateposts of K-GECO1 are colored in pink. (**B**) Systematic insertion scanning strategy by placing a linker at the mStayGold(J) gatepost - bulge region. (**C**) Alphafold3 model of insertion-tolerant i-mSG-v.1.0 with the inserted linker represented in red and positions of mutations represented in yellow.^24^ (**D**) The schematic representation of the initial prototype: HiCaRI v.0.0. Emission and excitation spectrum of HiCaRI v.0.0 in the presence and absence of 39 µM free Ca^2+^ (Δ*F*/*F*_min_ = −0.32). Excitation and emission maxima are at 499 nm and 510 nm, respectively. (**E**) Lineage of mSG variants generated in this work.

### Development of a mStayGold-based GECI (HiCaRI)

Using the insertion-tolerant i-mSG-v.1.0 as a starting point, we next aimed to construct a functional mStayGold-based GECI by inserting the CaM and ckkap Ca^2+^-binding domain from K-GECO1.^21^ Specifically, the flexible 8 amino acid linker sequence in i-mSG-v.1.0 (GGTG | GSGG) was split into four residue subsets, and a CaM and ckkap Ca^2+^-binding domain was inserted in between these linkers. While the initial prototype retained detectable fluorescence, it showed no Ca^2+^- dependent response. Subsequent linker optimization yielded a first mStayGold-GECI prototype (HiCARI v.0.0) that exhibited a clear Ca^2+^-dependent fluorescence change (Δ*F*/*F*_min_ = 0.33) (**Figure 1D**). However, during efforts to further improve fluorescence intensity and Ca^2+^ sensitivity through random mutagenesis, we identified a major limitation of this construct: a prolonged maturation time, which substantially hindered efficient screening and selection. Hence, we first prioritized improving maturation kinetics by selecting variants that exhibited faster fluorescence than the template construct. Through this selection process, we identified both direct- and inverse-response variants with Δ*F*/*F*_min_ of 0.24 and −0.32, respectively, with improved maturation time (**Figure 1E**). Based on performance and robustness during screening, we selected the inverse-response variant as the foundation for subsequent optimization. Eight additional rounds of directed evolution on this scaffold ultimately yielded HiCaRI, which exhibits a large inverse Ca^2+^-dependent fluorescence response (Δ*F*/*F*_min_ = −15) while preserving spectral properties of the original mStayGold(J).

The purified HiCaRI shows an inverse fluorescence response with a dynamic range *(ΔF*/*F*_min_) of −15 and high Ca^2+^ binding affinity (*K*_d_ = 38 nM) (**Figure 2A, B**). HiCaRI’s excitation and emission peaks are at 499 and 510 nm, respectively, consistent with the mStayGold(J) fluorescent protein (**Figure 2A**).^18^ In the presence of Ca^2+^, the chromophore p*K*_a_ increases from 6.0 to 8.2, and a distinct absorbance peak at 405 nm appeared, corresponding to the protonated (phenol) chromophore. This shift indicates that Ca^2+^ binding alters the chromophore protonation equilibrium, consistent with GCaMP-type sensors (**Figure 2B, C**).^25,26^ Additionally, the Ca^2+^- bound and Ca^2+^-free states exhibited quantum yields of 0.94 and 0.88 with extinction coefficients of 43,000 and 4,400 M^−1^ cm^−1^, respectively. The quantum yields of HiCaRI are among the highest reported for GFP-based GECIs, including the latest jGCaMP8s and mNG-GECO.^9,27^ Although molecular brightness in the Ca^2+^-unbound bright state is 3.4 times lower than mStayGold(J) and 1.8 times lower than mNG-GECO1 (Ca^2+^-bound, bright state), it is comparable to jGCaMP8s (Ca^2+^-bound, bright-state), with brightness approximately 1.3 times higher than that of the indicator (**Table 1**).

**Figure 2.**
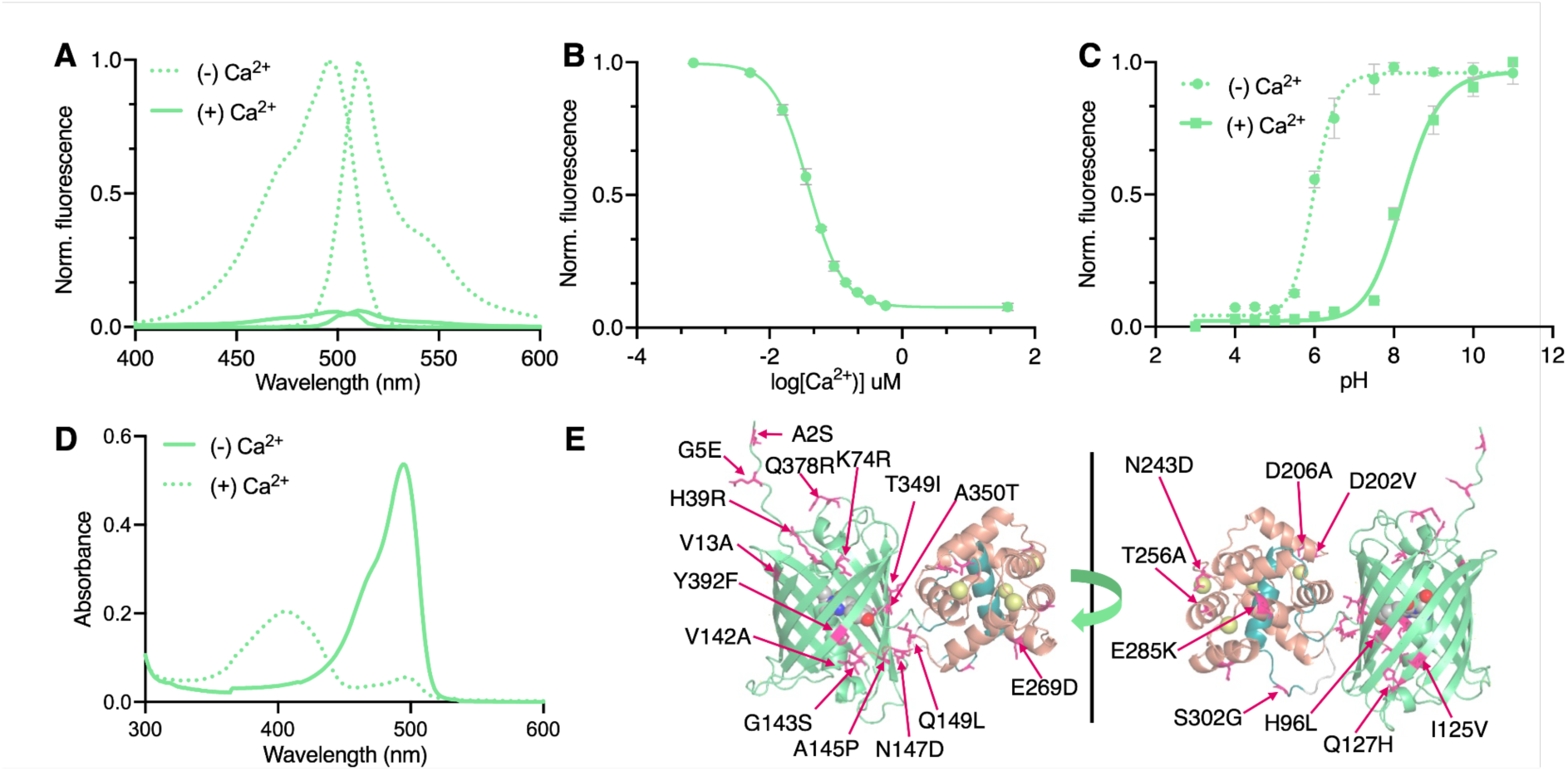
*In vitro* characterization of HiCaRI (**A**) Excitation and emission spectrum of purified HiCaRI protein in the presence (39 µM) and absence of free Ca^2+^. (**B**) Ca^2+^ titration of HiCaRI. *n* = 3 technical replicates (mean ± s.d.). (**C**) pH titration curves of HiCaRI with or without 39 µM Ca^2+^. *n* = 3 technical replicates (mean ± s.d.). (**D**) Absorbance spectra of HiCaRI in the presence (39 µM) and absence of Ca^2+^. (**E**) Alphafold3 predicted structure of HiCaRI, with mutated positions indicated.^24^

**Table 1.**
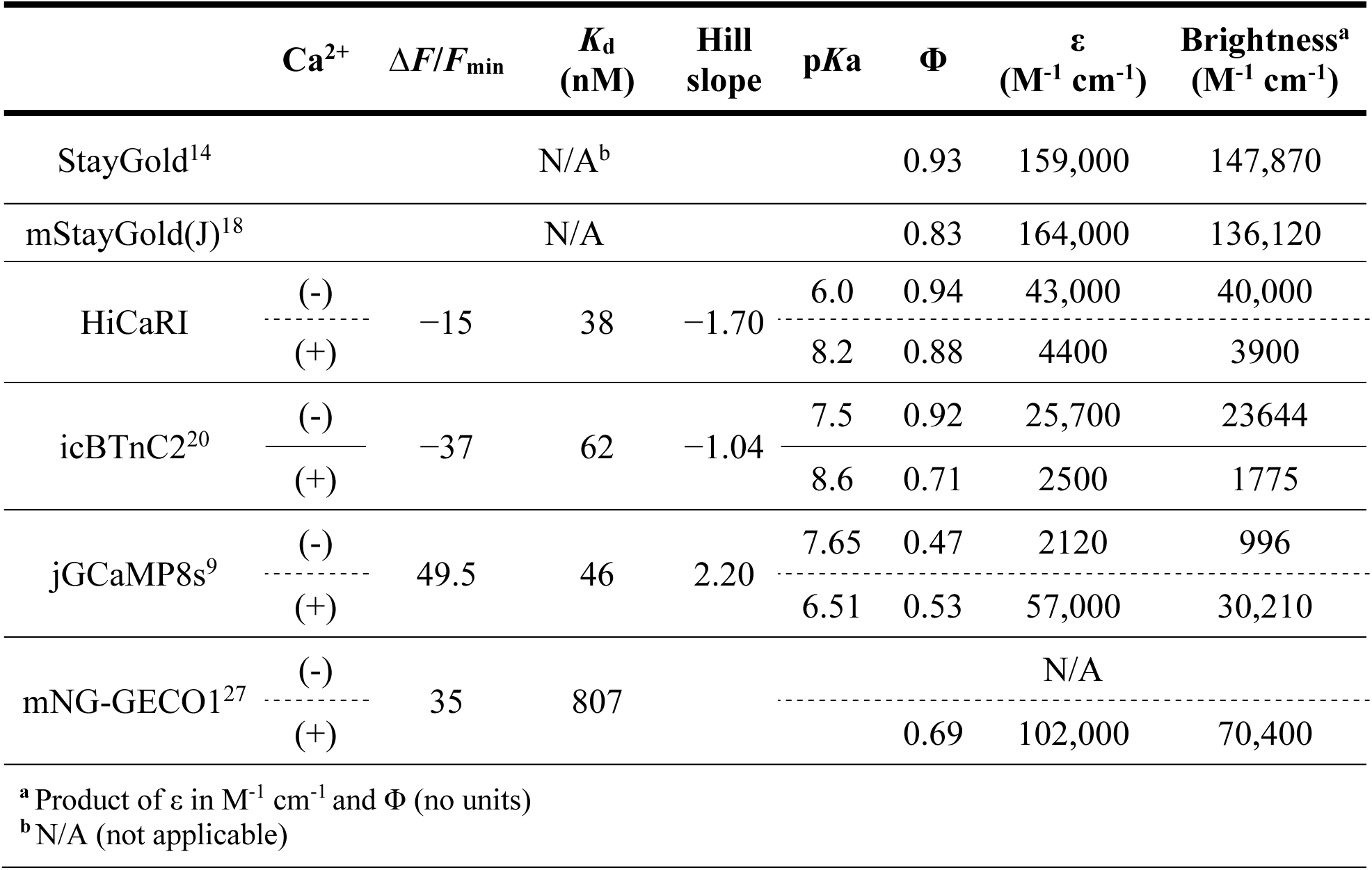
Characterization of HiCaRI and comparison against icBTnC2, jGCaMP8s, and mNG-GECO1.

The early rounds of directed evolution did not yield variants with |*ΔF*|/*F*_min_ greater than 1. However, we continued the evolution, first focusing on improving the expression and brightness (**Figure 2F**). In the 6^th^ and 7^th^ rounds, we identified variants containing mutations at sites 145, 147, and 149 at the linkers connecting i-mSG-v.1.0 and CaM, which exhibited |*ΔF*|/*F*_min_ greater than 1. A combination of these templates resulted in the current best variant, HiCaRI, with a *ΔF*/*F*_min_ of −15. Further investigation is necessary to identify the key mutations that increased sensitivity to Ca^2+^ and to improve affinity.

### Characterization of HiCaRI in mammalian cells

We validated the performance of HiCaRI in HeLa cells by adding histamine to induce intracellular Ca^2+^ oscillations, followed by EGTA/ionomycin treatment and subsequent Ca^2+^/ionomycin addition. HiCaRI was able to detect slight Ca^2+^ oscillations as well as reversible fluorescence change with the removal/addition of Ca^2+^ in HeLa cells (**Figure 3A, B, C)**. However, due to the high affinity of the current version, we observed an average *ΔF*/*F*_min_ of −0.46 ± 0.03 (n = 74 cells, mean ± s.e.m.), indicating a reduced fluorescence response compared with the ΔF/Fmin of −15 observed in purified proteins. The GECI was likely to be largely in the Ca^2+^-bound state at the resting intracellular Ca^2+^ concentration present in cells.

**Figure 3.**
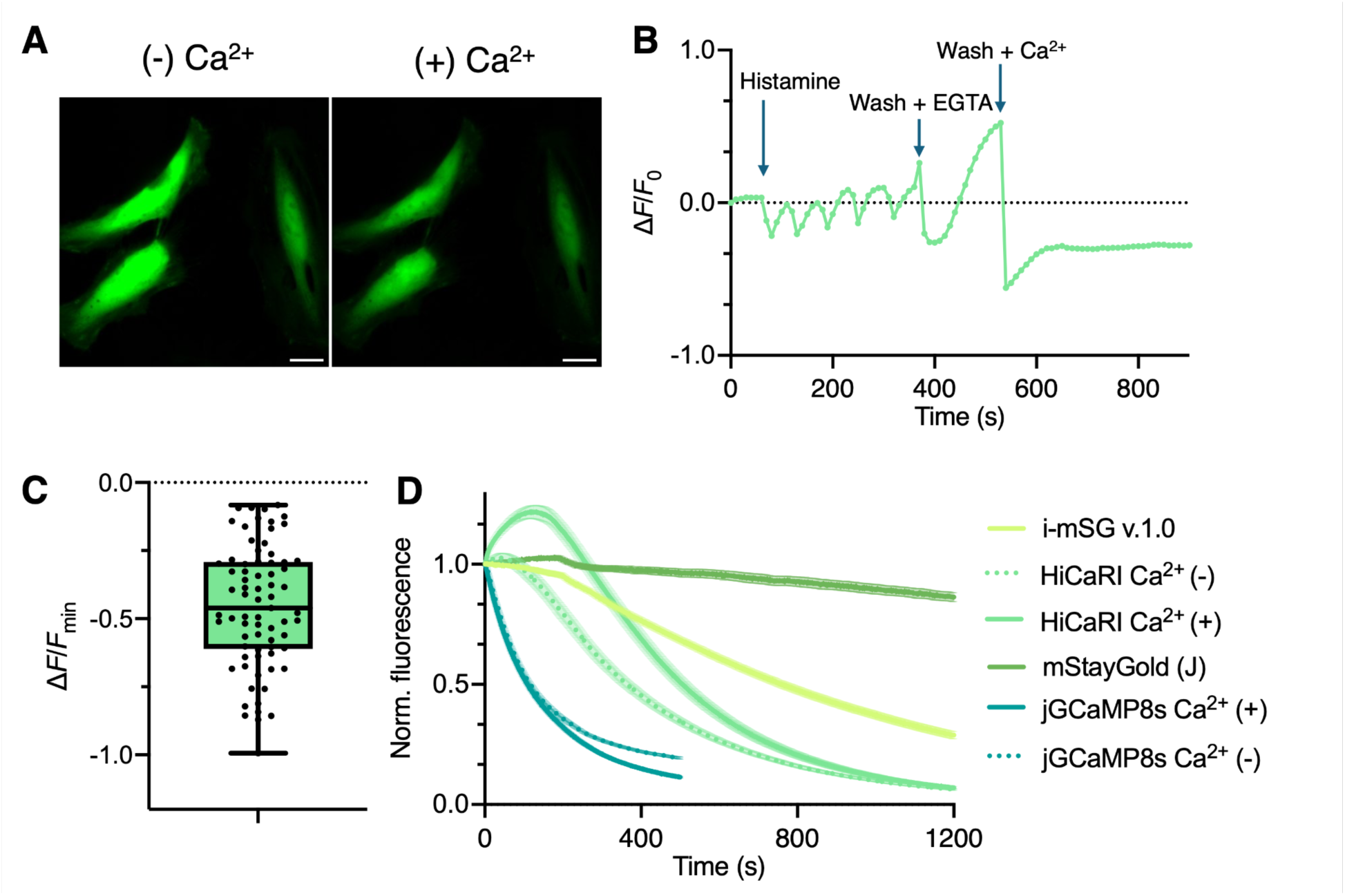
Mammalian cell-based characterization of HiCaRI (**A**) Image of HeLa cells expressing HiCaRI in the cytosol from a Ca^2+^-depleted (4 mM EGTA, 1.5 µM ionomycin, shown left) to a Ca^2+^ saturated state (10 mM CaCl_2_, 1.5 µM ionomycin, shown right). Scale bar = 20 µm. (**B**) Three-step Ca^2+^ fluorescence response of HiCaRI (5 µM histamine stimulation followed by 4 mM EGTA, 1.5 µM ionomycin, then 10 mM CaCl_2_, 1.5 µM ionomycin. Cells were seeded prior to EGTA and CaCl_2_ addition. (**C**) Boxplot of Δ*F*/*F*_min_ calculated from Ca^2+^ depletion to Ca^2+^ saturation over all sampled HeLa cells expressing HiCaRI in the cytosol (*n* = 74, *n* = 3 biological replicates, mean ± s.e.m). (**D**) Photostability comparison of H2B-HiCaRI and H2B-jGCaMP8s in the presence and absence of Ca^2+^. HeLa cells transfected with H2B-jGCaMP8s or H2B-HiCaRI were continuously illuminated (4.8 mW cm_−2_ at 470 ± 20 nm) for 500 or 1200 seconds. (*n* = 3 biological replicates, mean ± s.e.m.).

To evaluate the photostability of HiCaRI and compare it with the original mStayGold(J) and the jGCaMP8s biosensor, we created the corresponding histone 2B (H2B) fusion constructs and expressed them in cultured mammalian cells. H2B fusion provides nucleus-localization, limiting free diffusion through the cytosol and thereby facilitating photostability measurements.^28^ Photobleaching was assessed by continuously illuminating a fixed field of view for 500 or 1200 s (4.8 mW cm^−2^ at 470 nm). The fluorescence decay curves were fitted to a one-phase exponential decay model to calculate photobleaching half-lives (*t*_1/2_).

Consistent with previous reports, the control H2B-mStayGold(J) exhibited very high photostability, where more than 70% of the initial fluorescence was retained at 1200 s, indicating a photobleaching half-time exceeding 1200 s.^18^ However, H2B-i-mSG-v.1.0 showed reduced photostability relative to the control H2B-mStayGold(J) with a photobleaching half-life of 930 ± 71 s. Despite its reduced photostability, the photobleaching half-lives of H2B-HiCaRI (Ca^2+^-free, 330 ± 15 s; Ca^2+^-bound, 640 ± 83 s) were 3.9 and 6.5 times longer than H2B-jGCaMP8s in Ca^2+^-free and Ca^2+^-bound conditions, respectively (Ca^2+^-free, 85 ± 2.3 s; Ca^2+^-bound, 98 ± 1.4 s) (**Table 2**, **Figure 3D**).

**Table 2:**
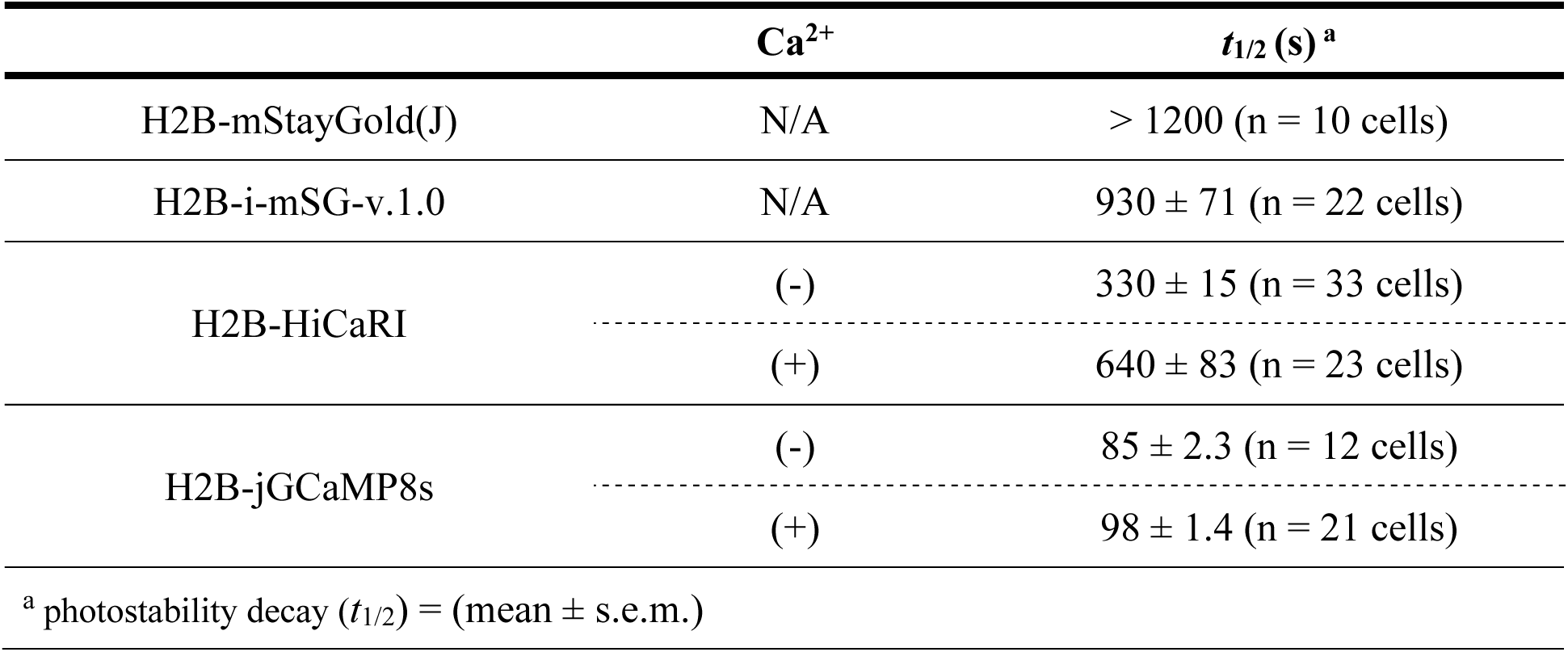
The photostability comparison.

## Discussion

In this work, we have demonstrated that a monomeric version of StayGold, a newly discovered green fluorescent protein with exceptional photostability and brightness, can be engineered to be a GECI through extensive stepwise engineering. The strategy we took was to first engineer an insertion-tolerant mStayGold(J) variant through a systematic insertion-site screening with a flexible linker as a placeholder to be later replaced with the Ca^2+^ sensing domain. Even though the initial prototype had less than 0.1% fluorescence intensity compared to the original mStayGold(J) (when expressed in *E. coli*), four rounds of directed evolution recovered much of the fluorescence and resulted in the insertion-tolerant i-mSG-v.1.0. We then aimed to engineer an initial prototype of a mSG-based GECI by replacing the flexible linker of i-mSG-v.1.0 scaffold with a CaM and ckkap Ca^2+^ sensing domain from K-GECO1. The initial prototype exhibited very slow chromophore maturation, which hindered the variant screening efficiency. Hence, we first focused on improving the maturation time and screened error-prone PCR libraries to identify variants with faster chromophore maturation. During the initial 4 rounds of directed evolution, we discovered direct- and inverse-response sensor prototypes with Δ*F*/*F*_min_ of 0.24 and −0.32, respectively. We first focused on evolving the inverse-response sensor prototype because we hypothesized that, due to the rigid structure of mStayGold(J), the conformational change upon Ca^2+^ binding would likely disrupt the structure, resulting in fluorescence quenching.

Eight additional rounds of directed evolution on the inverse sensor prototype produced HiCaRI with a Δ*F*/*F*_min_ of −15 upon Ca^2+^ binding, and a molecular brightness of 40,000 (Ca^2+^- unbound, bright-state), which is 1.3 times higher than jGCaMP8s (Ca^2+^-bound, bright-state). We then validated the response in HeLa cells. Due to the high Ca^2+^ affinity (*K*_d_ = 38 nM), HiCaRI, partially saturated at the resting intracellular Ca^2+^ concentration, which likely accounts for the average Δ*F*/*F*_min_ of −0.46 ± 0.03 observed in HeLa cells. Thus, HiCaRI may not be ideal for imaging applications requiring a *K*_d_ of several hundred nanomolar, suggesting that affinity tuning will be necessary for broader utility. Nevertheless, the current first-generation high-affinity variant would still be useful for detecting Ca^2+^ dynamics in cells with low-nanomolar resting Ca^2+^ levels and small transient signals.^29^

The evolution toward improved expression, brightness, and dynamic range in inverse-response sensors seems to compromise the exceptional photostability of the original mStayGold(J). Under continuous illumination in H2B-fused constructs, the original H2B-mStayGold(J) retained 70% of its initial fluorescence even after 1200 s of illumination. In contrast, the insertion-tolerant H2B-i-mSG-v.1.0 reached 50% of its initial fluorescence within 930 ± 71 s, indicating the reduced photostability compared to the original mStayGold(J). Nevertheless, H2B-HiCaRI exhibited the photobleaching half-lives of 330 ± 15 s (Ca^2+^-free) and 640 ± 83 s (Ca^2+^-bound), that is, 3.9 and 6.5 times longer than H2B-jGCaMP8s, respectively, suggesting the potential for a more photostable single FP-based GECI. This trade-off likely arises from the nature of directed evolution, which relies on random mutagenesis and high-throughput screening to improve a specific property of the biosensor. In our case, the insertion and the introduction of mutations increasing the fluorescence response (Δ*F*/*F*_min_) to Ca^2+^ perturbed structural features critical for maintaining photostability, which can be further explored by obtaining the crystal structure of our sensor construct. Future optimization will therefore require explicit selection for photostability, as well as detailed investigation of the mutations responsible for making mStayGold high in photostability. Recent advances in machine learning and high-throughput sequencing may further facilitate directed evolution by enabling a more comprehensive mapping of sequence-function relationships within a large mutant library.^30,31^ Implementing such approaches in directed evolution workflows for biosensor optimization can enable simultaneous optimization of multiple properties to minimize undesirable trade-offs between properties (i.e. Ca^2+^-dependent fluorescence change and photostability).^32,33^ Another design strategy we are pursuing is to circularly permutate our HiCaRI, repositioning the Ca^2+^ sensing domains (CaM and ckkap) at the new N- and C-termini. Such a design potentially allows greater Ca^2+^-dependent conformational changes to influence the chromophore, thereby increasing fluorescence response. This design has been widely adopted in high-performance GECIs and may further improve sensor performance, offering an alternative route to improved signal transduction while retaining the advantageous photophysical properties of the parent fluorescent protein.^5,34–36^ We are also continuing efforts to develop a direct-response HiCaRI.

In summary, we have demonstrated that a member of the new class of highly photostable mStayGold variants can be engineered into a high-performance functional GECI that exhibits greater photostability than a recent-generation GFP-based GCaMP biosensor. Our results suggest that mStayGold-based biosensors could offer a means of addressing the longstanding constraint of limited photostability, potentially expanding the palette of fluorescent protein-based biosensors suited for long-term, high light intensity, imaging applications.

## Methods

### General methods and materials

mStayGold(J) construct was kindly donated by Professor Miyawaki from RIKEN. Phusion high-fidelity DNA polymerase (Thermo Fisher Scientific) was used for standard gene amplification, while Taq DNA polymerase (New England Biolabs) was used for error-prone PCR. Site-directed mutagenesis was performed using the QuikChange mutagenesis kit (Agilent Technologies). Restriction endonucleases, Rapid DNA Ligation Kit, and GeneJET Miniprep Kits were purchased from Thermo Fisher Scientific. PCR and restriction digest products were purified using agarose gel electrophoresis and the GeneJET Gel Extraction Kit (Thermo Fisher Scientific). The HeLa cell line was purchased from ATCC (#CCL-2). Absorbance spectra were measured using the Shimadzu UV1800 spectrometer. Fluorescence emission spectra were recorded on a Spark plate reader (Tecan). DNA sequences were analyzed by Fasmac Co., Ltd.

### Protein purification and *in vitro* characterization

The constructs cloned into pBAD-HisB with an N-terminal 6×His tag were expressed in the *E. coli* strain DH10B (Thermo Fisher Scientific). A single colony from freshly transformed E. coli was inoculated into two tubes of 5 mL of Terrific Broth (TB, 1 μL/mL ampicillin) at 37 °C overnight. The seed culture was then added to fresh TB (2% Luria-Bertani Broth supplemented with an additional 1.4% tryptone, 0.7% yeast extract, 54 mM K_2_PO_4_, 16 mM KH_2_PO_4_, 0.8% glycerol) with ampicillin (100 μL/100 mL of TB) and incubated at 37 °C until OD600 reached the exponential growth phase (OD > 0.6). Once the OD600 reached 0.6, L-arabinose (200 µL/100 mL TB) was added to induce expression for another 16 h at room temperature. Bacteria were then harvested and lysed using a sonicator (Branson). After centrifugation (12,000 × g, 20 min), the supernatant was purified with Ni^2+^-NTA affinity agarose beads (G-Biosciences) on a column (Thermo Fisher ScientificTM PierceTM Centrifuge Columns 10 mL). The eluted sample was further concentrated and desalted with a 30 kDa Amicon Ultra−15 Centrifugal Filter. The protein concentration was determined by absorbance at 280 nm (Nanodrop, Thermo Scientific). Absorbance spectra were measured using the Shimadzu UV1800 spectrometer. To perform pH titrations, protein solutions were diluted into buffers (pH from 3 to 11) containing 50 mM citrate, 50 mM Tris, 50 mM glycine, 100 mM NaCl, and either 2 mM CaCl_2_ or 2 mM EGTA.^9^ A sigmoidal binding function was then fitted to fluorescence intensities as a function of pH to determine the p*K*_a_ for Ca^2+^-positive and Ca^2+^-negative conditions. For Ca^2+^ titration, buffers were prepared by mixing a zero free Ca^2+^ buffer (30 mM MOPS, 10 mM EGTA, 100 mM KCl, pH 7.2) and 39 μM free Ca^2+^ (+) buffer (30 mM MOPS, 10 mM CaEGTA, 100 mM KCl, pH 7.2) to provide Ca^2+^ concentrations ranging from 0 to 39 μM at 25 °C. ^32^ Fluorescence intensities were plotted against Ca^2+^ concentrations and fitted by a sigmoidal binding function in GraphPad Prism to determine the Hill coefficient and apparent *K*_d_ value. All fluorescence measurements using a plate reader were performed using the same conditions (emission mode, λ_ex_ = 460 ± 20 nm, λ_em_ = 510 ± 20 nm).

### Characterization of HiCaRI in HeLa cells

The constructed genes in pBAD (mStayGold(J), i-mSG-v.1.0, HiCaRI, jGCaMP) were digested with XhoI and HindIII, and ligated into pcDNA3.1 (Thermo Fisher Scientific) vectors for cytoplasmic expression in mammalian cells. H2B constructs used for photostability experiments were cloned in mPapaya1-H2B-6 backbone using Gibson Assembly (Addgene plasmid #56651). HeLa cells were maintained in Dulbecco’s modified Eagle medium (DMEM high glucose; Nacalai Tesque) supplemented with 10% fetal bovine serum (FBS; Sigma-Aldrich) and 1% penicillin-streptomycin (Nacalai Tesque) at 37 °C and 5% CO_2_. Cells were seeded in 35-mm glass-bottom cell culture dishes (Iwaki) and transiently transfected with constructed plasmids using polyethyleneimine (Polysciences) before imaging. After 48-72 hours, transfected cells were imaged using an IX83 wide-field fluorescence microscopy (Olympus) equipped with a pE-300 LED light source (CoolLED), a 40× objective lens (numerical aperture (NA) = 1.3; oil), an ImagEM X2 EM-CCD camera (Hamamatsu), Cellsens software (Olympus), and an STR stage incubator (Tokai Hit). The filter sets used for live-cell imaging had the following specifications: excitation 470/20 nm, dichroic mirror 490 nm dclp, and emission 518/45 nm. Fluorescence images were analyzed with ImageJ software (National Institutes of Health).

For the cell-based Ca^2+^ imaging, adherent HeLa cells seeded onto a glass-bottom dish were washed twice with Hank’s balanced salt solution (+) (HBSS (+); Nacalai Tesque, 09735-75), and the buffer was exchanged to 900 μL HBSS (+) supplemented with 10 mM HEPES (Nacalai Tesque, 17557-94) just before imaging. Then histamine (Wako, 087-03533, 50 μM final concentration) was added one minute after the start of imaging. At the indicated time, imaging was paused, and the dish was washed twice with HBSS without Ca^2+^ (HBSS (-)) (Nacalai Tesque, 672 17461-05), and 1 mL of HBSS (-) was added. Then, EGTA/ionomycin (4 mM/1.5 μM final concentration) was added to quench the Ca^2+^-dependent oscillation. Imaging was stopped again at the indicated time, and the dish was washed twice more with HBSS (+), and 1 mL of HBSS (+) was added. Right before resuming, Ca^2+^/ionomycin (10 mM/1.5 μM final concentration) was immediately added to saturate the indicator.^33^ The photostability experiment was performed by transfecting HeLa cells with H2B-mSG, H2B-i-mSG-v.1.0, H2B-jGCaMP8s, or H2B-HiCaRI. 48-72 hours after transfection, cells were washed with HBSS(-) or HBSS(+) buffers. Then, the cells were incubated with either EGTA/ionomycin (4 mM/1.5 μM final concentration) in HBSS(-) or Ca^2+^/ionomycin (10 mM/1.5 μM final concentration) in HBSS(+) for at least 5 min. The fluorescence of the sensors localized in the nucleus was recorded under continuous illumination (4.8 mW cm^−2^ at 470 nm). Fluorescence images were analyzed using ImageJ software (National Institutes of Health).

## Data Availability Statement

The authors declare that the data supporting the findings of this study are available within the paper and its Supplementary Information files. Reasonable requests for reagents or raw data files in another format should be directed to a corresponding author.

## Acknowledgments

The authors thank Professor Atsushi Miyawaki from RIKEN for kindly sharing mStayGold(J) plasmid. This research was supported by the Japan Society for the Promotion of Science (JSPS) (21H00273 and 23H02101 to T.T., and 24H00489 and 24H02267 to R.E.C.) and an award from Japan Science and Technology Agency (JST) CREST (JPMJCR25T3) and the Mitsubishi Foundation. I.M. is supported by JST SPRING (JPMJSP2108) and the Forefront Physics and Mathematics Program to Drive Transformation (FoPM), a World-leading Innovative Graduate Study (WINGS) Program of The University of Tokyo.

## Author Contributions

I.M. designed and performed all experiments unless otherwise noted. K.K.T., K.T-Y., and R.E.C. guided the design of HiCaRI and assisted with *in vitro* experiments. K.T-Y. assisted and supervised in cell imaging experiments. R.E.C. conceived the project. K.K.T., K.T-Y., T.T., and R.E.C. supervised the research. I.M. wrote the first draft of the paper, with guidance and editing from K.K.T., K.T-Y., and R.E.C. All authors read and approved the final manuscript.

## Notes

The authors declare no competing financial interest.

