## Supplementary material for "A StayGold-based calcium ion indicator"

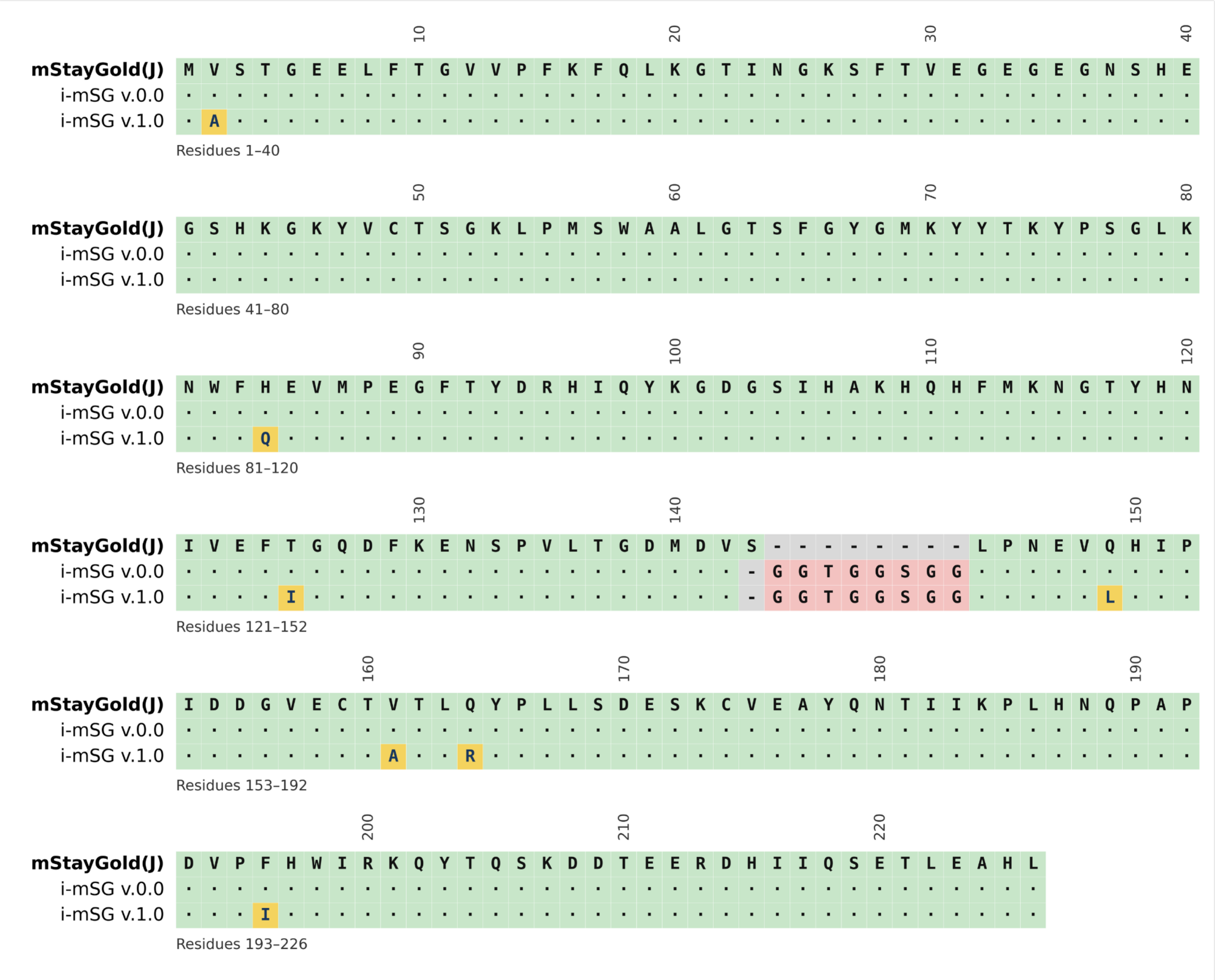


**Supplementary Figure 1. Sequence alignment of mStayGold(J) and i-mSG v.1.0.** Yellow indicates a mutation, and pink residues are the mock linker used to construct the insertion-tolerant i-mSG v.1.0.

**Supplementary Table 1. DNA and protein sequences of variants in this study**

**Sequence (5' to 3')**

**>i-mSG_v.0.0_DNA**
atggtctctactggagaggaacttttcacgggggtcgttccgttcaaatttcaactcaagggaacgatcaatggtaaatccttcacagtggaagg
agaaggtgaagggaactcacacgagggaagccataaagggaagtacgtctgcacttctgggaaacttccgatgagctgggcagcactggg
aactagcttcggatatgggatgaaatactatacgaaatacccttccgggcttaagaactggtttcatgaagtgatgccggagggatttacttacga
taggcatattcaatacaagggtgatggatcaattcatgcgaaacatcagcatttcatgaagaacggaacttatcataacattgttgagtttaccgg
gcaagactttaaggaaaattctcctgttttgacaggagacatggacgtaggcggcaccggcggcagcggcggccttcctaacgaggtacagc
atatcccaatcgacgatggcgtcgagtgcactgttactctgcagtacccccctgttgtctgatgagtccaagtgtgtagaagcatatcaaaacacc
atcatcaaacccctgcataaccaaccagctcccgatgttccttttcattggatcagaaagcaatacacacaaagtaaggatgacacagaaga
acgagaccatattattcagtccgagactttggaggcgcacctt

**>i-mSG_v.0.0_protein**
MVSTGEELFTGVVPFKFQLKGTINGKSFTVEGEGEGNSHEGSHKGKYVCTSGKLPMSWAALGTSFG
YGMKYYTKYPSGLKNWFHEVMPEGFTYDRHIQYKGDGSIHAKHQHFMKNGTYHNIVEFTGQDFKEN
SPVLTGDMDVGGTGGSGGLPNEVQHIPIDDGVECTVTLQYPLLSDESKCVEAYQNTIIKPLHNQPAPD
VPFHWIRKQYTQSKDDTEERDHIIQSETLEAHL

---

**>i-mSG_v.1.0_DNA**
atggcctctactggagaggaacttttcacgggggtcgttccgttcaagtttcaactcaagggaacgatcaatggtaaatccttcacagtggaagg
agaaggtgaagggaactcacacgagggaagccataaagggaagtacgtctgcacttctgggaaacttccgatgagctgggcagcactggg
aactagcttcggatatgggatgaaatactatacgaaatacccttccgggcttaagaactggtttcaagaagtgatgccggagggatttacttacg
ataggcatattcaatacaagggtgatggatcaattcatgcgaaacatcagcatttcatgaagaacggaacttatcataacattgttgagtttatcg
ggcaagactttaaggaaaattctcctgttttgacgggagacatggacgtaggcggcaccggcggcagcggcggccttcctaacgaggtactg
catatcccaatcgacgatggcgtcgagtgcactgctactctgcggtacccccctgttgtctgacgagtccaagtgtgtagaagcatatcaaaacac
catcatcaaacccctgcataaccaaccagctcccgatgttcctattcattggatcagaaagcaatacacacaaagtaaggatgacacagaag
aacgagaccatattattcagtccgagactttggaggcgcacctt

**>i-mSG_v.1.0_protein**
MASTGEELFTGVVPFKFQLKGTINGKSFTVEGEGEGNSHEGSHKGKYVCTSGKLPMSWAALGTSFG
YGMKYYTKYPSGLKNWFQEVMPEGFTYDRHIQYKGDGSIHAKHQHFMKNGTYHNIVEFIGQDFKEN
SPVLTGDMDVGGTGGSGGLPNEVLHIPIDDGVECTATLRYPLLSDESKCVEAYQNTIIKPLHNQPAPD
VPIHWIRKQYTQSKDDTEERDHIIQSETLEAHL

---

**>HiCaRI v.0.0_DNA**
atggcctctactggagaggaacttttcacgggggtcgttccgttcaagtttcaactcaagggaacgatcaatggtaaatccttcacagtggaagg
agaaggtgaagggaactcacacgagggaagccataaagggaagtacgtctgcacttctgggaaacttccgatgagctgggcagcactggg
aactagcttcggatatgggatgaaatactatacgagatacccttccgggcttaagaactggtttcaagaagtgatgccggagggatttacttacg
ataggcatattcaatacaagggtgatggatcaattcatgcgaaacatcagcatttcatgaagaacggaacttatcataacattgttgagtttatcg
ggcaagactttaaggaaaattctcctgttctgacgggagacatggacgcaggcgaggccagtaatggccaactgactgaagagcagatcgc
agaatttaaagaggctttctccctatttgacaaggacggggatgggacgataacaaccaaggagctggggacggtgatgcggtctctggggc
agaaccccacagaagcggagctgcaggacatgatcaatgaagtagatgccgacggtgacggcacattcgacttccctgagttcctgacgat
gatggcaagaaaaatgagttatagagtcactgaagaggaaattagagaagcgttccgcgtgtttgataaggacgtcaatggctacatcggcg
cagcagagcttcgccacgtgatgacagaccttggagagaagttaacagatgaggaggttgatgaaatgatcagggtagcagacatcgatgg
ggatggtcaggtaaactacgaagagtttgtacaaatgatgacagcgaagggtggcggaggttctatgggcagcgtgaagctgatccccagcc
tgaccaccgtgatcctcgtgaagtccatgctgcggaagcggagcttcggcaaccccttcaagtatggcggccttcctaacgaggtactgcatat
cccaatcgacgatggcgtcgagtgcattgccactctgcggtacccccctgttgtctgacgagtccaagtgtgtagaagcatatcaaaacaccatc
atcaaacccctgcataaccaaccagctcccgatgttcctattcattggatcagaaagcaatacacacaaagtaaggatgacacagaagaac
gagaccatattattcagtccgagactttggaggcgcacctt

**>HiCaRI v.0.0_protein**
MASTGEELFTGVVPFKFQLKGTINGKSFTVEGEGEGNSHEGSHKGKYVCTSGKLPMSWAALGTSFG
YGMKYYTRYPSGLKNWFQEVMPEGFTYDRHIQYKGDGSIHAKHQHFMKNGTYHNIVEFIGQDFKEN

SPVLTGDMDAGEASNGQLTEEQIAEFKEAFSLFDKDGDGTITTKELGTVMRSLGQNPTEAELQDMINE
VDADGDGTFDFPEFLTMMARKMSYRVTEEEIREAFRVFDKDVNGYIGAAELRHVMTDLGEKLTDEEV
DEMIRVADIDGDGQVNYEEFVQMMTAKGGGGSMGSVKLIPSLTTVILVKSMLRKRSFGNPFKYGGLP
NEVLHIPIDDGVECIATLRYPLLSDESKCVEAYQNTIIKPLHNQPAPDVPIHWIRKQYTQSKDDTEERDH
IIQSETLEAHL

---

**>HiCaRI v.0.1_DNA**
atggcctctactggagaggaacttttcacgggggtcgttccgttcaagtttcaactcaagggaacgatcaatggtaaatccttcacagtggaagg
agaaggtgaagggaactcacacgagggaagccataaagggaagtacgtctgcacttctgggaaacttccgatgagctgggcagcactggg
aactagcttcggatatgggatgaaatactatacgaaatacccttccgggcttaagaactggtttcaagaagtgatgccggagggatttacttacg
ataggcatattcaatacaagggtgatggatcaattcatgcgaaacatcagcatttcatgaagaacggaacttatcataacattgttgagtttatcg
ggcaagactttaaggaaaattctcctgttttgacgggagacatggacgtaggcgaggccagtaatggccaactgactgaagagcagatcgca
gaatttaaagaggctttctccctatttgacaaggacggggatgggacgataacaaccaaggagctggggacggtgatgcggtctctggggca
gaaccccacagaagcggagctgcaggacatgatcaatgaagtagatgccgacggtgacggcacattcgacttccctgagttcctgacgatg
atggcaagaaaaatgagttatagagtcactgaagaggaaattagagaagcgttccgcgtgtttgataaggacgtcaatggctacatcggcgc
agcagagcttcgccacgtgatgacagaccttggagagaagttaacagatgaggaggttgatgaaatgatcagggtagcagacatcgatggg
gatggtcaggtaaactacgaagagtttgtacaaatgatgacagcgaagggtggcggaggttctatgggcagcgtgaagctgatccccagcct
gaccaccgtgatcctcgtgaagtccatgctgcggaagcggagcttcggcaaccccttcaagtatggcggccttcctaacgaggtactgcatatc
ccaatcgacgatggcgtcgagtgcactgccactctgcggtacccccctgttgtctgacgagtccaagtgtgtagaagcatatcaaaacaccatca
tcaaacccctgcataaccaaccagctcccgatgttcctattcattggatcagaaagcaatacacacaaagtaaggatgacacagaagaacg
agaccatattattcagtccgagactttggaggcgcacctt

**>HiCaRI v.0.1_protein**
MASTGEELFTGVVPFKFQLKGTINGKSFTVEGEGEGNSHEGSHKGKYVCTSGKLPMSWAALGTSFG
YGMKYYTKYPSGLKNWFQEVMPEGFTYDRHIQYKGDGSIHAKHQHFMKNGTYHNIVEFIGQDFKEN
SPVLTGDMDVGEASNGQLTEEQIAEFKEAFSLFDKDGDGTITTKELGTVMRSLGQNPTEAELQDMINE
VDADGDGTFDFPEFLTMMARKMSYRVTEEEIREAFRVFDKDVNGYIGAAELRHVMTDLGEKLTDEEV
DEMIRVADIDGDGQVNYEEFVQMMTAKGGGGSMGSVKLIPSLTTVILVKSMLRKRSFGNPFKYGGLP
NEVLHIPIDDGVECTATLRYPLLSDESKCVEAYQNTIIKPLHNQPAPDVPIHWIRKQYTQSKDDTEERD
HIIQSETLEAHL

---

**>HiCaRI_DNA**
atgtcctctactgaagaggaacttttcacaggggtcgctccgttcaagtttcaactcaagggaacgatcaatggtaaatccttcacagtggaagg
agaaggcgaagggaactcacgcgagggaagccataaagggaagtacgtctgcacttctgggaaacttccgatgagctgggcagcactgg
gaactagcttcggatatgggatgaaatactatacgaggtacccttccgggcttaagaactggtttcaagaagtgatgccggagggatttacttac
gataggcttattcaatacaagggtgatggatcaattcatgcgaaacatcagcatttcatgaagaacggaacttatcataacattgttgagtttgtcg
ggcatgactttaaggaaaattctcctgttctgacgggagacatggacgcaagcgagcccagtgatggcctactgactgaagagcagatcgca
gaatttaaagaggctttctccctatttgacaaggacggggatgggacgataacaaccaaggagctggggacggtgatgcggtctctggggca
gaaccccacagaagcggagctgcaggacatgatcaatgaagtagttgccgacggtgccggcacattcgacttccctgagttcctgacgatga
tggcaagaaaaatgagttatagagtcactgaagaggaaattagagaagcgttccgcgtgtttgataaggacgtcgatggctacatcggcgca
gcagagcttcgccacgtgatggcagaccttggagagaagttaacagatgaggaggttgatgatatgatcagggtagcagacatcgatgggg
atggtcaggtaaactacaaagagtttgtacaaatgatgacagcgaagggaggtggaggttctatgggcggcgtgaagctgatccccagcctg
accaccgtgatcctcgtgaagtccatgctgcggaagcgtagcttcggcaaccccttcaagtacggcggcctccctaacgaggtactgcatatcc
caatcgacgatggcgtcgagtgcattaccactctgcggtacccccctgttgtctgacgagtccaagtgtgtagaagcatatcaaaacaccatcatc
aaacccctgcataaccgaccagctcccgatgttcctattcattggatcaggaagcaattcacacaaagtaaggatgacacagaagaacgag
accatattattcagtccgagactttggaggctcacctttaa

**>HiCaRI_protein**
MSSTEEELFTGVAPFKFQLKGTINGKSFTVEGEGEGNSREGSHKGKYVCTSGKLPMSWAALGTSFG
YGMKYYTRYPSGLKNWFQEVMPEGFTYDRLIQYKGDGSIHAKHQHFMKNGTYHNIVEFVGHDFKEN
SPVLTGDMDASEPSDGLLTEEQIAEFKEAFSLFDKDGDGTITTKELGTVMRSLGQNPTEAELQDMINE
VVADGAGTFDFPEFLTMMARKMSYRVTEEEIREAFRVFDKDVDGYIGAAELRHVMADLGEKLTDEEV
DDMIRVADIDGDGQVNYKEFVQMMTAKGGGGSMGGVKLIPSLTTVILVKSMLRKRSFGNPFKYGGLP
NEVLHIPIDDGVECITTLRYPLLSDESKCVEAYQNTIIKPLHNRPAPDVPIHWIRKQFTQSKDDTEERDHI
IQSETLEAHL
